# Re-evaluating donor-cell identity after hiPSC-RGC transplantation in mouse models of optic neuropathy

**DOI:** 10.64898/2026.11.11.766031

**Authors:** Vrathasha Vrathasha, Sergei Nikonov, Brent A. Bell, Kimberly E. Dine, Yingrui Liu, Reas S. Khan, Kenneth S. Shindler, Ahmara G. Ross, Venkata R. M. Chavali, Joan M. O’Brien

## Abstract

Glaucoma is a neurodegenerative disease characterized by permanent visual field loss caused by the death of retinal ganglion cells (RGCs). Glaucoma is the leading cause of irreversible blindness worldwide. We investigated the potential of transplanting human induced pluripotent stem cell-derived RGCs (hiPSC-RGCs) in two mouse models of optic nerve degeneration: *N*-Methyl-D-Aspartate (NMDA) and optic nerve crush (ONC), and characterized their ability to survive, integrate, and form synaptic connections in a degenerated retina. Follow-up studies, including immunohistochemistry and patch-clamping experiments, one year after intravitreal injections, determined the location, identity, and function of the donor GFP^+ve^ fluorescent cells within the mouse retinas and their similarity to native RGCs. Upon closer inspection, some GFP^+ve^ cells detected in a few retinal sections did not stain for the human-specific marker and were detected in retinal layers outside the expected ganglion cell layer (GCL)/ inner nuclear layer (INL) regions. These findings highlight complications in the interpretation of RGC transplantation studies.

## INTRODUCTION

Glaucoma is a chronic neurodegenerative disease characterized by progressive structural and functional decline of the optic nerve, resulting in permanent loss of vision. The clinical characteristics of glaucoma have been attributed to the death of retinal ganglion cells (RGCs).^1^ RGCs, like most neuronal cells, are post-mitotic and hence lack self-regenerative capacity. The loss of RGCs disrupts the transmission of visual stimuli from photoreceptors in the eye to the brain, leading to irreversible visual field loss.^2^ In 2022, about 4.22 million people in the US were estimated to be suffering from glaucoma^3^, and according to the CDC glaucoma programs, this number is projected to increase to 6.3 million cases by the year 2050.

The insidious progression of glaucoma results in delayed diagnosis, when treatment is often less effective.^4^ As there are no remedies available once RGC death has occurred, cell-mediated therapy has been proposed as an alternative therapy for glaucoma.^5^ Optimization of this approach could potentially restore vision in glaucoma patients. This methodology involves transplanting RGCs derived from human induced pluripotent stem cells (hiPSCs)^6,7^ into the vitreous cavity of optic neuropathy murine eyes to replace the function of dead ganglion cells.

Previously, we showed that green fluorescent protein (GFP)-labeled donor hiPSC-RGCs survived in wild-type C57BL/6J mice for up to 5-months post transplantation. Furthermore, the transplanted hiPSC-RGCs were well-integrated, expressed RGC- and human-specific markers, and were shown to be electrophysiologically active within the mouse retina.^8^ In this study, we evaluated the feasibility of cell replacement therapy in two mechanistically distinct murine models of RGC loss, namely, *N*-Methyl-D-Aspartate (NMDA)-induced excitotoxicity and optic nerve crush (ONC)-induced optic neuropathy. Survival of the transplanted hiPSC-RGCs was assessed by fundus photography and histological studies one year after transplantation. Two-photon live imaging of retinal explants showed that surviving GFP^+ve^ cells had RGC-like morphology and exhibited electrophysiological properties comparable to those of native RGCs.

Studies of retinal cell transplantation have raised concerns about intercellular cytoplasmic material transfer between donor and host photoreceptor cells.^9-13^ Specifically, Zhang et al., 2023, reported that RGC xenografts occasionally transferred donor-derived cytoplasmic, nuclear, and mitochondrial proteins to host Müller glia. This process, although rare, was greatly facilitated by the disruption of the inner limiting membrane (ILM).^14^ Therefore, careful assessment of the identity of the GFP-labeled cells in our study revealed that in some sections of the mouse retina, cells with robust GFP expression did not stain for human nuclear antigen. Transfer/leakage of fluorescent proteins from donor cells to host cells can lead to false positives, confounding the analysis of these types of cell transplantation studies.

## RESULTS

### Fluorescently labeled cells were detected in the murine retina of the optic neuropathy eyes

To provide a pre-clinical framework necessary to advance RGC replacement therapy as a treatment for glaucoma, studies were conducted in murine models of RGC loss. We selected two mouse models of optic neuropathy: NMDA-induced excitotoxicity and ONC.^15-19^ In the NMDA mouse model, H&E and fluorescence immunohistochemistry (IHC) staining of Brn3^+ve^ cells showed a dose-dependent reduction in RGC numbers across increasing NMDA concentrations (1.5 mM, 3 mM, and 6 mM) (**Figure S1**). At 1.5 mM, NMDA caused significant RGC death without detectable changes in other neuro-retinal layers; thus, this dose was used for all our subsequent NMDA injections in C57BL/6 mice.^20^ Whereas ONC-induced optic neuropathy involves an acute mechanical injury directly to the optic nerve, causing RGC death.

For our transplantation studies in NMDA- and ONC-optic neuropathy mouse models, hiPSC-RGCs were fluorescently labeled by transduction with an AAV2.7m8-*SNCG*-eGFP vector, in which eGFP expression was driven by RGC-specific gamma Synuclein (*SNCG*) promoter. Four weeks after the NMDA- and ONC-induced RGC loss, 2 µL of 1 X 10^6^ of hiPSC-RGCs were injected into the vitreous space of a C57BL/6 mouse’s eye (8-week-old, *n*=10 per mouse model). The contralateral eye received 2 µL of 1X PBS and served as the experimental control. Confocal scanning laser ophthalmoscopy (cSLO) detected GFP^+ve^ cells at 2, 4, and 6 weeks after transplantation in 9 of 10 NMDA-treated mice and 5 of 10 ONC-injured mice (**Figure 1**). The mice were periodically monitored for any signs of complications. Inflammatory responses were elevated in the retinas of mice that underwent optic nerve injury, and survival of donor hiPSC-RGCs was attenuated in ONC mice. Thus, all subsequent data reported in this manuscript are primarily from the NMDA-treated mice unless mentioned otherwise.

**Figure 1:**
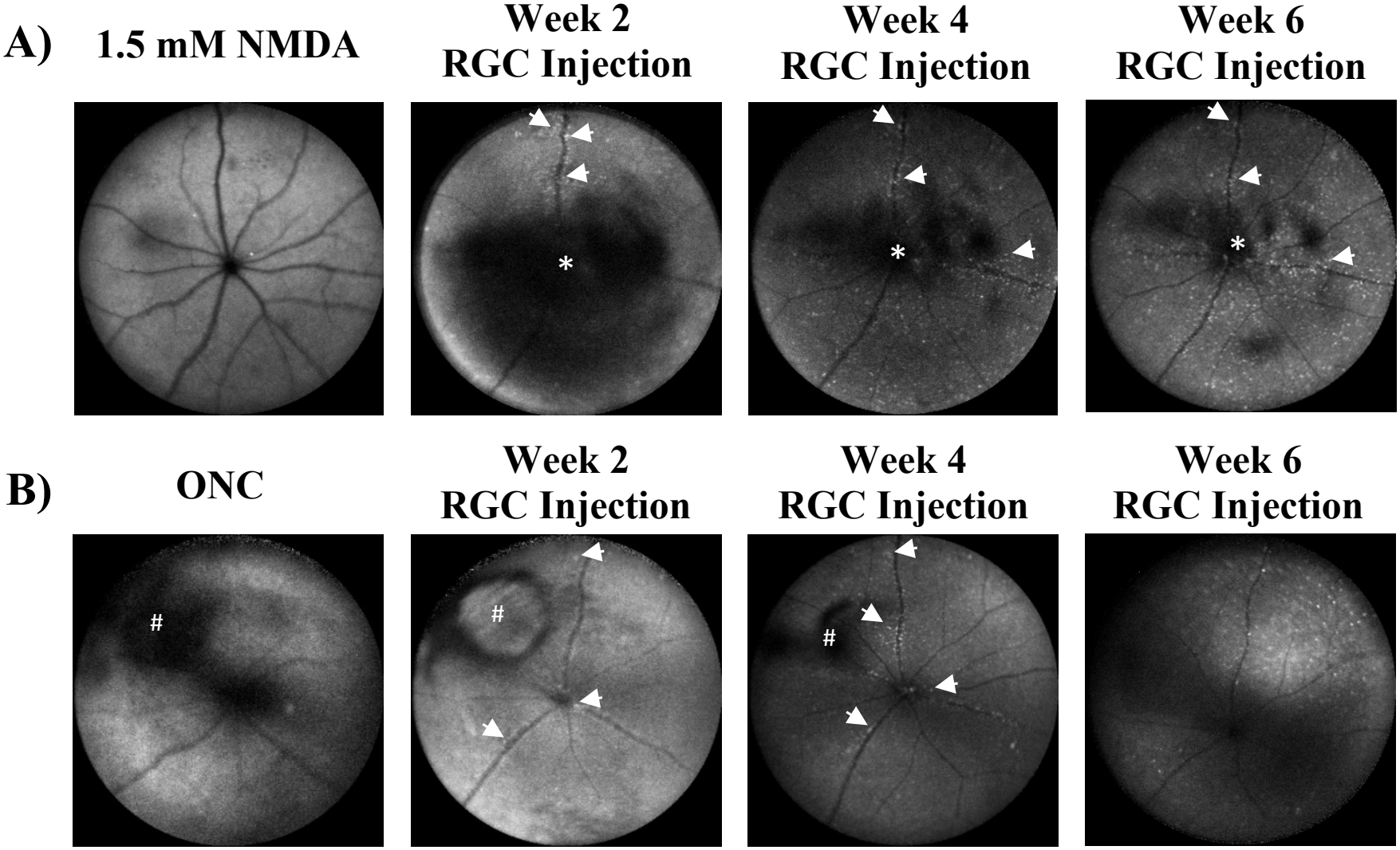
Fundus images of GFP^+ve^ fluorescent cells in NMDA- and ONC-mouse models. Purified hiPSC-RGCs transduced with the *SNCG*-eGFP vector were intravitreally injected into NMDA-treated and ONC-injured C57BL/6 mice. Images of fluorescent cells were taken using cSLO at 2, 4, and 6 weeks in A) 1.5 mM NMDA-injected mice and B) ONC-injured mice. In mice subjected to ONC, we detected an elevated inflammatory response, as visible in week 6 after RGC injection. * Indicative of post-injection damage resolving over time. # Indicative of damage caused by optic nerve crush.

### GFP^+ve^ cells detected in the murine retina exhibited RGC and synaptic markers

One year after hiPSC-RGC transplantation into mice that received 1.5 mM of NMDA, healthy eyes without any abnormalities were enucleated for retinal whole-mount IHC (*n*=6), and Z-stack images were taken using confocal microscopy. We documented and compared morphological and histological changes in the enucleated NMDA-treated eyes to determine the location and the integration of the graft hiPSC-RGCs in the mouse retina.

Flattened retinas were counterstained for RGC-specific markers such as BRN3, RBPMS, and TUJ1, including synaptic markers like VGLUT2, PSD95, and syntaxin-1. *SNCG*-eGFP^+ve^ cells detected in the mouse retinas expressed BRN3 and RBPMS (**Figure 2A-B**), and GFP^+ve^ neurites co-localized with the TUJ1 marker (**Figure 2C**). Furthermore, the GFP^+ve^ cells expressed syntaxin-1 in the host retina (**Figure 2D**). *SNCG*-eGFP^+ve^ cells are shown here to express PSD95 protein in the neurite terminal, which has the characteristic punctate pattern (**Figure 2E**). Co-staining with VGLUT2 showed that *SNCG*-eGFP^+ve^ cells expressed this vesicular transporter (**Figure 2F**).

**Figure 2:**
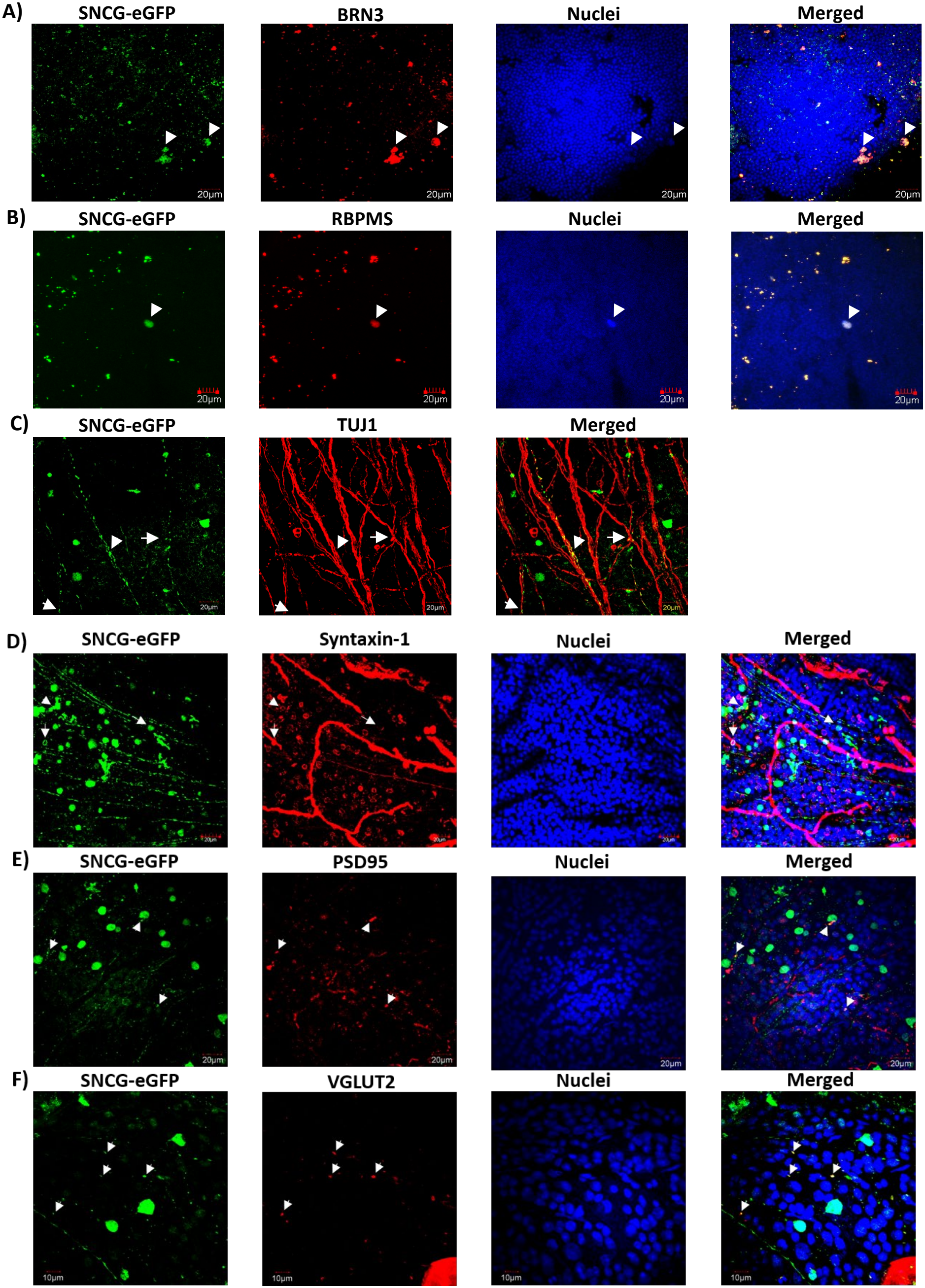
Characterization of *SNCG*-GFP^+ve^ cells in the mouse retina. (A–F) Flattened whole-mount images of GFP^+ve^ cells co-stained for the following markers: (A) BRN3, (B) RBPMS, (C) TUJ1, (D) Syntaxin-1, (E) PSD95, and (F) VGLUT2.

### The GFP-labeled cells were determined to be electrophysiologically active within the mouse retina

We evaluated the functional analysis of the GFP^+ve^ cells in the murine retina using electrophysiological recordings from retinal flat mounts one year after transplantation. Fluorescent cells were identified by two-photon imaging and targeted for recording in cell-attached/voltage-clamp and/or whole-cell/current-clamp configurations under infrared illumination. Each recording was followed by two-photon imaging of the recorded cell. Robust light-evoked responses were observed in 8 of 11 recorded cells.

Representative images and light responses are shown in **Figures 3** and **S2. Figure 3** illustrates the morphology and light response of cell 1, whereas **Figure S2** summarizes all eight light-responsive cells, including six from NMDA-treated and two recorded from ONC-injured retinas.

**Figure 3:**
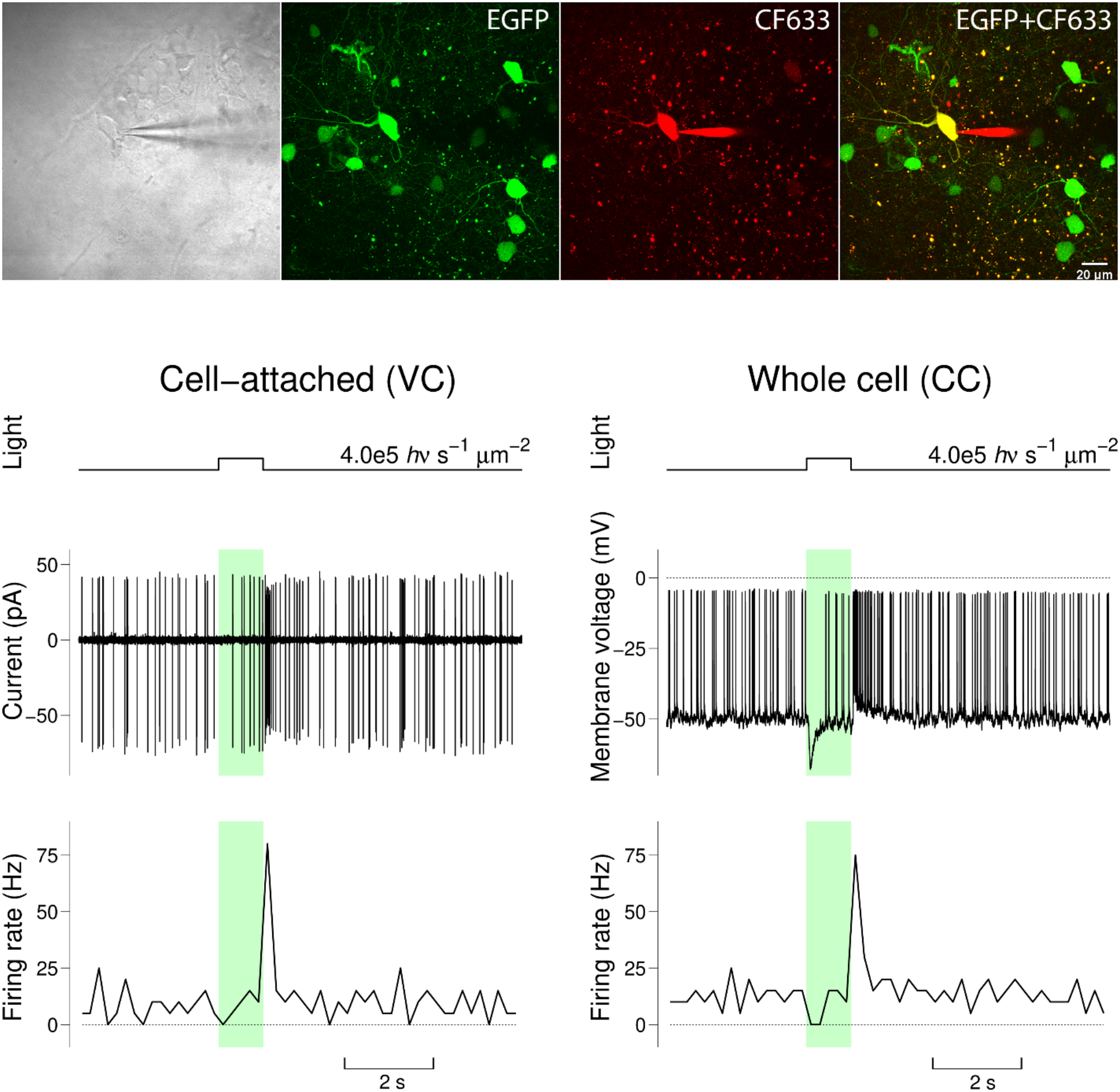
Light responses of a GFP^+ve^ cell integrated into the mouse retina. Top panels show transmitted-light images together with green fluorescence (eGFP expressed by the recorded cell), red fluorescence (CF633 included in the pipette solution), and the merged green/red channels acquired using two-photon microscopy. Traces below illustrate light responses of the targeted cell recorded in cell-attached/voltage-clamp (left column) and whole-cell/current-clamp (right column) configurations. For each recording configuration, the top trace indicates the timing of light stimulation, the middle trace shows either the high-pass-filtered pipette current (cell-attached/voltage-clamp mode) or membrane potential (whole-cell/current-clamp mode), and the bottom trace shows the spike firing rate calculated using 200-ms time bins. Sharp vertical deflections in the current and voltage traces represent action potential spikes. Colored bars indicate periods of light stimulation. Light intensities are indicated on the corresponding graphs. Membrane potential was not corrected for the liquid junction potential; corrected values would be approximately 16 mV more negative.

The recorded GFP^+ve^ cells exhibited the major classes of light responses expected in wild-type RGCs. Four cells (cells 2, 3, 4, and 8) exhibited ON responses, two cells (cells 1 and 5) exhibited OFF responses, and two cells (cells 6 and 7) exhibited ON-OFF responses. Three additional cells, all from NMDA-treated retinas, are not included in the figures. Two displayed unusually high spontaneous firing rates and less clearly defined light responses, likely due to damage incurred during patching. The third exhibited delayed OFF responses to approximately 60% of light stimuli despite maintaining a healthy resting membrane potential (∼-60 mV, uncorrected for liquid junction potential) and a low spontaneous firing rate.

Optokinetic responses (OKR) were measured every three months for up to a year to evaluate visual acuity after hiPSC-RGC injections in NMDA-treated eyes (**Figure S3**).^21^ OKR was measured in both eyes: the right eyes that were treated with 1.5 mM of NMDA and hiPSC-RGCs, and the contralateral eyes injected with 1.5 mM of NMDA and 1X PBS. OKR measurements were taken until the mice were sacrificed one year after hiPSC-RGC injections (*n*=9). ANOVA did not show a significant difference in OKR scores in eyes injected with hiPSC-RGCs across different time points when compared with the 1X PBS-injected control eyes.

### Origin of the GFP^+ve^ cells observed in the murine retina

Retinal cryosections and whole-mount retinas were co-stained with human-specific markers, including human nuclear antigen (HuNu) and Ku80, to confirm the origin of the GFP^+ve^ cells. In certain retinal sections, *SNCG*-eGFP^+ve^ cells expressed both HuNu and Ku80 (**Figure 4A-C**). However, some retinal sections also showed many *SNCG*-eGFP^+ve^ cells that did not stain for the HuNu marker (**Figure 4D-E**).

Furthermore, videos of the retina generated from two-photon z-stacks taken from the ganglion cell layer (GCL) toward the photoreceptor layers show that some non-RGC cells exhibited green fluorescence. As expected, the majority of the GFP^+ve^ cells were located in the GCL. However, some GFP^+ve^ cells were also detected within the inner nuclear layer (INL). The retina shown in **Video 1** exhibited the highest proportion of GFP^+ve^ cells outside the GCL. Cell counts from this retina identified approximately 73 GFP^+ve^ RGCs, 20 presumptive bipolar or amacrine cells located in the inner INL and exhibiting faint arborization, and 11 presumptive horizontal cells located in the outer INL and characterized by brighter, thicker processes.

**Figure 4:**
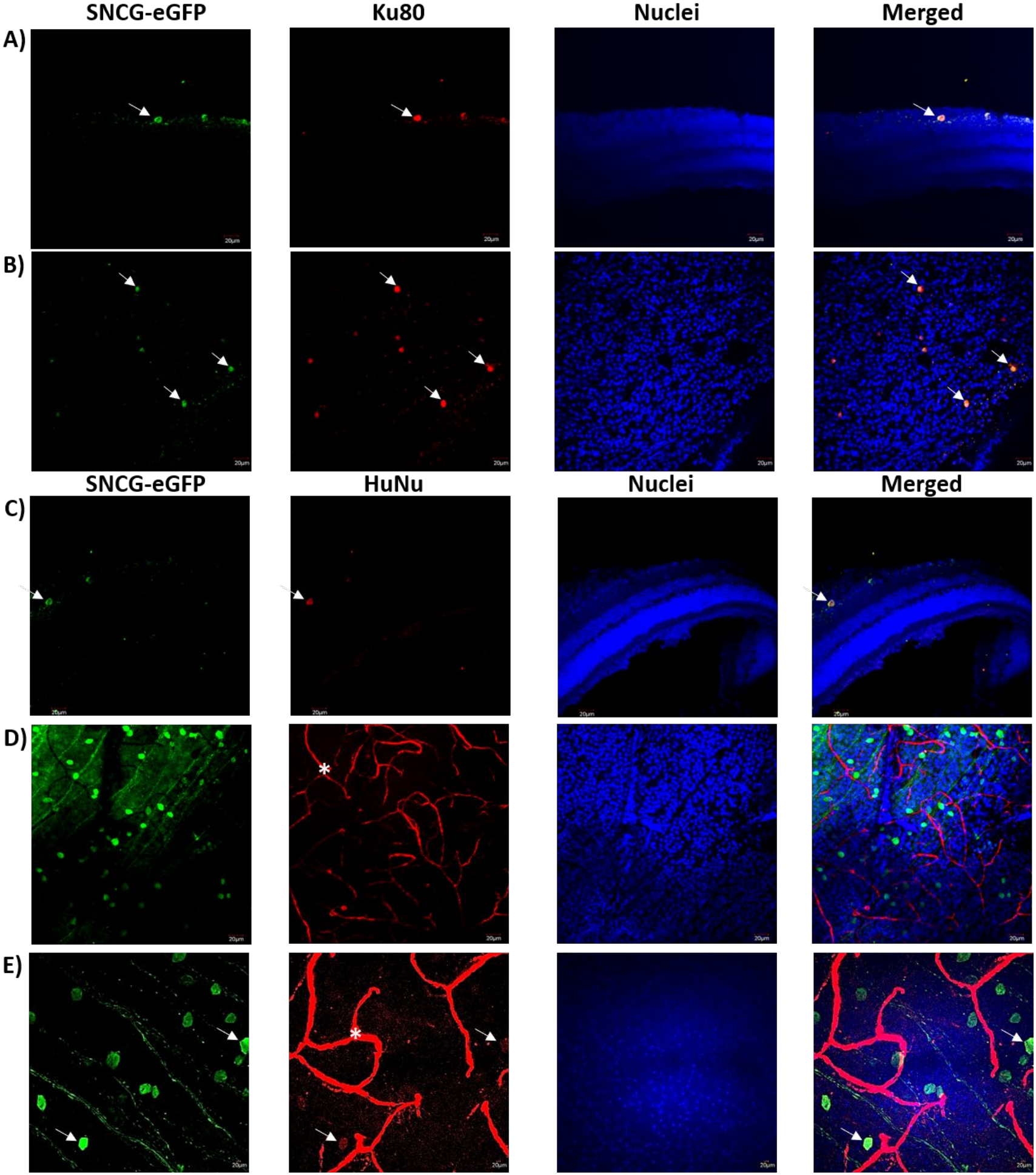
Immunostaining of the *SNCG*-eGFP^+ve^ cells in the mouse retina. (A, C) Retinal cryosections and (B, D, E) flattened whole-mount images of eGFP^+ve^ cells co-stained with human-specific markers (A-B) Ku80 and (C-E) human nuclear antigen (HuNu) to confirm the origin of the fluorescent cells. * Since the human nuclear antigen antibody was raised in mice, some cross-reactivity with the murine retina was observed.

## DISCUSSION

Glaucoma is the most common type of optic neuropathy and is characterized by progressive loss of RGCs and degeneration of the optic nerve. RGCs are the retina’s only projection neurons, collecting visual information from other retinal neurons and transmitting it via the optic nerve to visual targets in the brain. Thus, RGC death leads to irreversible vision loss. Several groups, including our team, have proposed using RGC cell replacement therapy as an alternative treatment to restore vision in glaucoma patients.^14,22-27^

In this study, we evaluated the long-term survival and characteristics of fluorescently labeled hiPSC-RGCs after transplantation in two mouse models of RGC loss. GFP^+ve^ cells remained detectable for up to one year following transplantation and exhibited RGC-associated morphological, molecular, and electrophysiological properties. However, a subset of fluorescent cells lacked human-specific markers and were observed outside the expected retinal layers, raising questions about cellular identity.

We have previously reported the survival and functionality of donor hiPSC-RGCs after intravitreal injection in wild-type C57BL/6J mice.^8^ The cells were pre-labelled with AAV2.7m8-*SNCG*-eGFP to express eGFP. Transplanted hiPSC-RGCs survived and integrated within the ganglion cell layer in 15 of 16 mice. At 2-5 months post-transplantation, graft-derived fluorescent cells stained positive for RGC-specific markers, synaptic markers, human nuclear antigen, and Ku80. Two-photon imaging and patch-clamp electrophysiology of live retinal explants confirmed the morphology and function of eGFP^+ve^ hiPSC-RGCs, and donor axons extended into the optic nerve, where their expression colocalized with a neurofilament marker.

The present study extended this work to two optic neuropathy models in order to evaluate donor cell survival, integration, and synaptic connectivity in a degenerating retina. In the NMDA model, intravitreally delivered NMDA binds to glutamate receptors on neuronal cells, causing calcium influx and dose-dependent apoptosis of RGCs.^15-17^ In the ONC model, injury is inflicted unilaterally on the orbital portion of the optic nerve, producing progressive optic nerve degeneration, changes at the optic nerve head, and RGC death.^18^ Following crush injury, RGC survival in wild-type mice falls from 47% at day 7 to 27% at day 14 relative to controls.^19^ However, unilateral ONC also induces the expression of pro- and anti-inflammatory molecules and gliosis markers.^28^ As observed at week 6 post hiPSC-RGC injections, inflammatory responses were elevated in our ONC mice, and this may be one factor affecting overall donor cell survival (**Figure 1B**).

We examined donor hiPSC-RGCs for up to one year after transplantation using fundus imaging, OKR, and histological retinal analyses. To our knowledge, this study represents the longest reported follow-up period for RGC transplantation. Fluorescent cells were detected within the retina and, based on IHC, exhibited overlap with established RGC and synaptic markers. Live two-photon imaging of retinal explants further revealed cellular morphologies consistent with RGC-like features, while electrophysiological recordings provided evidence of neuronal activity in the fluorescently labeled population. OKR measurements, however, showed no significant improvement in hiPSC-RGC-injected eyes relative to PBS-injected contralateral eyes at any time point, and no functional benefit was therefore demonstrated at the behavioral level. Despite these findings, we could not definitively attribute all fluorescent cells to the donor human cells, as some lacked detectable human-specific marker expression and localized to retinal layers beyond the anticipated GCL/INL. One way to distinguish mouse and human retinal cells would be by their nuclear morphologies, as mouse nuclei are small and have prominent nucleoli, whereas human nuclei show homogeneous DAPI staining.^14^ In our study, we have included an image of a flat-mount retina transplanted with hiPSC-RGCs, showcasing a mixture of *SNCG*-eGFP^+ve^ cells with nuclei stained with homogeneous DAPI and nuclei with prominent nucleoli (**Figure S4**). This unexpected labeling pattern introduces uncertainty regarding the cellular identity and origin of the fluorescent population.

The detection of GFP-positive cells that are not positive for human nuclear antigen may be explained by at least two potential mechanisms. First, fluorescent material may have been transferred from the transplanted human RGCs to endogenous mouse retinal cells through intercellular material exchange. Alternatively, residual AAV2 carried over with the transplantation cell preparation may have resulted in the unintended transduction of host mouse cells. Even though we detected the presence of Iba1, indicative of microglia, we consider phagocytic uptake of dying transplanted hiPSC-RGCs by inflammatory cells to be less likely, as the GFP signal in these non-human cells appeared as diffuse, uniform cytoplasmic fluorescence rather than the punctate pattern typically associated with material contained within phagolysosomes (**Figure 4**). In addition, given the one-year interval following transplantation, we would expect cellular debris from nonviable GFP^+ve^ hiPSC-RGCs to have been cleared by this time.

Intercellular material transfer is well documented between donor and host rod photoreceptors.^9-13^ However, it was unclear until recently if donor RGCs also engage in intercellular material transfer. Zhang et al., 2023 noted that in conditions where the ILM is removed, RGC xenografts do take part in rare donor-derived protein transfer to host Müller glia.^14^ What seems to be apparent is that material transfer between donor and host cells is dramatically enhanced in degenerative ocular conditions. Similarly, in our studies of hiPSC-RGC transplantation in wild-type mice, we did not observe any intercellular transfer of cytoplasmic proteins between donor and host cells. Granted, we cannot definitively rule out that no material transfer occurred in wild-type eyes, as we were limited by the number of retinal samples available for staining with species-specific markers. However, in the case of NMDA-mediated RGC loss, some of the fluorescent-positive cells detected in the mouse retina did not stain for a human-specific marker. NMDA is known to cause substantial inner-retinal injury, including loss of RGCs and amacrine cells, and induces strong reactive Müller gliosis.^29,30^ Zhang et al., 2023 reported that Müller glial cells were recipients of donor-derived cytoplasmic proteins.^14^ We detected the presence of Müller glial and microglial cells in the retinas, as evidenced by staining for GFAP and Iba1 markers, respectively (**Figure S5A-B**). However, their expression did not colocalize with *SNCG*-GFP^+ve^ cells. Since we did not stain for other Müller glial-specific markers such as vimentin or nestin, we cannot rule out this possibility. However, expression of *SNCG-*GFP^+ve^ cells did colocalize with bipolar neurons, as seen by PKC-α staining (**Figure S5C**), consistent with the laminar position and morphology of the GFP^+ve^ INL cells observed in **Video 1**.

With respect to the possibility of residual AAV carryover leading to transduction of mouse cells after intravitreal injections, it is reported that even a tiny presence of AAV2 can efficiently transduce murine RGCs and express GFP for over a year.^31-33^ If the donor hiPSC-RGCs were not sufficiently washed before intravitreal injection into the murine eye, this would explain why non-human cells were GFP^+ve^. AAV2 binding studies have shown that substantial cell-associated viral material, even after washing, does not remove virions associated with the cell surface or internalized into the cell.^34^ Incomplete removal of vector before intravitreal injection would therefore account for GFP expression in host murine neurons. Furthermore, the 7m8 capsid was engineered for enhanced penetration of the ILM from the vitreous^35^; studies have shown that optic nerve injury and NMDA-treated models alter retinal architecture, causing AAV2 to penetrate the inner/GCL retina, including Müller cells and bipolar neurons.^36,37^

In conclusion, following the successful transplantation of RGCs in wild-type mice, we sought to replicate our previous transplantation studies in two optic neuropathy mouse models characterized by NMDA- and ONC-mediated RGC loss. We evaluated the survival of transplanted hiPSC-RGCs using fundus photography and histological analyses for up to one year after transplantation. IHC demonstrated co-localization of green fluorescence with RGC-specific and synaptic markers. Furthermore, two-photon live imaging of retinal explants and electrophysiological analyses provided evidence supporting the morphology and functional properties of the fluorescently labeled cells that were similar to those of an expected RGC. However, a subset of fluorescent cells did not stain for the human-specific marker and were detected in retinal layers outside the expected GCL/INL regions, complicating the interpretation of our findings. These observations highlight a critical challenge in definitively establishing the identity and origin of fluorescently labeled cells following transplantation. The inability to conclusively distinguish transplanted cells from endogenous host-derived cells represents an important limitation in the interpretation of RGC transplantation studies and, more broadly, in stem cell-based transplantation research. Future retinal transplantation studies cannot rely solely on fluorescent reporter expression to establish donor-cell survival and integration.

## LIMITATIONS

Experiments that conclusively prove the identity of the GFP^+ve^ cells detected in the mouse retina by staining for AAV vector DNA, a species-specific marker, and an RGC-specific marker will be the subject of future studies. We would also perform sex-mismatched Y chromosome FISH or stain donor nuclei with a thymidine analog and the cytoplasm with lacZ or GFP; these experiments provide useful information on the identity, position, and morphology of the fluorescently labeled cells.^9^

Furthermore, in this study, we cannot conclusively determine whether the GFP-labeled cells subjected to patch-clamp experiments were human donor cells or mouse host cells. In future experiments, the cell bodies of patched cells could be collected for RNA analysis to confirm their species and cellular identity.

## EXPERIMENTAL MODEL AND SUBJECT DETAILS

### Human iPSC culture

Undifferentiated human-induced pluripotent stem cells (hiPSCs) were derived and characterized by DNA fingerprinting, karyotype analysis, qRT-PCR, and FACS, thereby providing a comprehensive assessment of iPSC characteristics.^38-40^ All human sample collection protocols were approved by the University of Pennsylvania and the Children’s Hospital of Philadelphia Human Subjects Research Institutional Review Board following the Declaration of Helsinki. All methods were performed per the relevant research guidelines and regulations of the University of Pennsylvania. Written informed consent was obtained from all human cell donors. The iPSCs were maintained in an iPSC medium: StemMACs iPSC-Brew XF (Miltenyi Biotec, catalog #: 130-107-086, North Rhine-Westphalia, Germany) containing 50X supplement (Miltenyi Biotec, catalog #: 130-107-087, North Rhine-Westphalia, Germany), and 1% Penn/Strep (Life Technologies, catalog #: 10378-016, Carlsbad, CA, USA).

### Retinal progenitor cell (RPC) generation and conditions

hiPSCs were seeded in a 6-well tissue culture dish (Corning, catalog # 229106, Corning, NY, USA) coated with 1:100 diluted growth factor reduced Matrigel Growth Factor (GFR; Corning, catalog #: 354230, Corning, NY, USA) and were maintained in an iPSC medium + 20 ng/mL of bFGF and 5 ng/mL of stable bFGF. Two days post-plating, media were collected, filtered, and either used directly or cryopreserved for later use. hiPSCs were maintained at 37 °C at 5% O2 and 5% CO2 until reaching 100% confluence then transferred to 37 °C, 5% CO2 overnight prior to induction. On day 0, iPSC media was changed to RPC induction media: DMEM/F12 (50:50; Corning, catalog #: 10-092-cm, Corning, NY, USA), 1% Penn/Strep (Life Technologies, catalog #: 10378-016, Carlsbad, CA, USA), 1% Glutamine MAX (Life Technologies, catalog #: 35050-061, Carlsbad, CA, USA), 1% NEAA (Life Technologies, catalog #: 11140-050, Carlsbad, CA, USA), 0.1 mM 2-ME (Life Technologies, catalog #: 21985-023, Carlsbad, CA, USA), 2% B27 supplement (w/o vitamin A; Life Technologies, catalog #: 12587-010, Carlsbad, CA, USA), 1% N2 supplement (Life Technologies, catalog #: 17502-048, Carlsbad, CA, USA), containing 2 μM XAV939 (X) (Wnt inhibitor; R&D Systems, catalog #: 3748, Minneapolis, MN, USA), 10 μM SB431542 (SB) (TGF-β inhibitor; R&D Systems, catalog #: 1614, Minneapolis, MN, USA), 100 nM LDN193189 (L) (BMP inhibitor; R&D Systems, catalog #: 6053, Minneapolis, MN, USA), 10 mM nicotinamide (Sigma-Aldrich, catalog #: N0636, St. Louis, MO, USA), and 10 ng/mL IGF1 (R&D Systems, catalog #: 291-G1, Minneapolis, MN, USA). Cultures were fed daily for 4 days. On day 4, culture media were exchanged with RPC induction media containing: 2 μM XAV939, 10 μM SB431542, 100 nM LDN193189, 10 ng/mL IGF1, and 10 ng/mL bFGF. Media were changed daily for 21 days.^7^

### Retinal ganglion cell (RGC) differentiation

Before RGC differentiation, the medium was changed daily using RGC induction media containing 250 ng/mL Shh (R&D Systems, catalog #: 8908-SH, Minneapolis, MN, USA), 100 ng/mL FGF8 (R&D Systems, catalog #: 423-F8, Minneapolis, MN, USA) for 2 days. For RGC differentiation on day 24, cells were manually crossed into small clusters with Leibovitz’s medium containing 34 μM D-Glucose (Research Products International, catalog #: G32045-500, Mt. Prospect, IL, USA) using the crosshatching technique as previously described (Espuny-Camacho, Michelsen et al. 2013), then re-plated at a density of 1.0 × 10^5^ cells/well of 6-well plate coated with 1:100 diluted growth factor reduced Matrigel Growth Factor in RGCs induction media containing: 100 ng/mL Follistatin 300 (R&D Systems, catalog #: 669-FO, Minneapolis, MN, USA), 0.5 μM Cyclopamine (R&D Systems, catalog #: 1623/1, Minneapolis, MN, USA), 3 μM DAPT (Stemgent, catalog #: 04-0041, Cambridge, MA, USA), and 4.2 µM Y-27632 dihydrochloride (ROCK inhibitor; R&D Systems, catalog #: 1254/1, Minneapolis, MN, USA). Media was changed 24 hours post-plating with RGC induction media containing: 100 ng/mL Follistatin 300 and 3 μM DAPT daily for 2 days. From day 27, media was changed to RGC induction media containing: 3 μM DAPT, 10 µM ROCK inhibitor (Y27632), 5 μM Forskolin (Selleckchem, catalog #: S2449, Houston, TX, USA), 400 μM N^6^,2′-O-Dibutyryladenosine 3′,5′-cyclic monophosphate sodium salt (cAMP; Sigma-Aldrich, catalog #: D0627, St. Louis, MO, USA), 40 ng/mL BDNF (R&D Systems, catalog #: 248-BDB, Minneapolis, MN, USA), 5 ng/mL NT4 (R&D systems, catalog #: 268-N4, Minneapolis, MN, USA), and 10 ng/mL CNTF (R&D systems, catalog #: 257-NT, Minneapolis, MN, USA). Medium was changed every 2–3 days until day 36. After maturation (D36), the medium was exchanged every 3–4 days with RGC induction media containing: 3 μM DAPT and 10 µM ROCK inhibitor (Y27632).^7^

### AAV vector design and production

AAV expression cassettes were flanked by the AAV2 inverted terminal repeats. A pro-viral plasmid driving expression of enhanced green fluorescent protein (eGFP)^41^ was driven by codon-optimized *SNCG* (gamma-synuclein promoter)^42^ and contains identical cis-regulatory elements. AAV2.7m8-SNCG.eGFP vector was generated and purified with a CsCl gradient by the CAROT research vector core at the University of Pennsylvania.^43^ hiPSC-RGCs (Day 40) were transduced with AAV2.7m8 virus-containing SNCG-eGFP vector at a concentration of 1.92 X 10^10^ vg/mL. hiPSC-RGCs expressed green fluorescence as early as 48 hours post-transduction; the cells were washed with 1X PBS three days after the virus was first applied, and media were changed at least two times before hiPSC-RGCs were intravitreally injected (∼Day 50) into the eye of C57BL/6J mice.

### Animal husbandry

Wild-type C57BL/6J mice were obtained from the Jackson Laboratory (Bar Harbor, ME, USA) and raised in a 12-h light/dark cycle. Animals were housed at the University of Pennsylvania in compliance with the ARVO Statement for the Use of Animals in Ophthalmic and Vision Research as well as with institutional and federal regulations.

## METHOD DETAILS

### Intravitreal injections of NMDA

1-month-old wild-type C57BL/6J mice were anesthetized with a mixture of xylazine and ketamine administered intraperitoneally, with 0.5% proparacaine eye drops applied topically. Powdered NMDA (≥98%, catalog #: M3262, Sigma-Aldrich, St. Louis, MO, USA) was dissolved in 0.1 mol/L phosphate-buffered saline (PBS) to obtain final concentrations of 1.5 mM, 3 mM, and 6 mM. For NMDA standardization studies, 1 µL of 1.5 mM, 3 mM, and 6 mM NMDA was injected into the vitreous space of both eyes (*n*=5/treatment group) using a 33-gauge 0.5” needle attached to a 5 μL Hamilton syringe (Hamilton Company, catalog #: 7803-05, Reno, NV, USA). Following the injections, Prednisolone 1%/ Gentamicin 0.3% ophthalmic ointment was applied topically to the eyes. The animals were placed on a heating pad until awake and monitored periodically for any signs of distress or complications.

For transplantation studies, 1 µL of 1.5 mM NMDA was injected into the vitreous space of both eyes (*n*=10) using a 33-gauge 0.5” needle attached to a 5 μL Hamilton syringe (Hamilton Company, catalog #: 7803-05, Reno, NV, USA). The final NMDA concentration in the eye is estimated to be ∼250 µM, 1/6th of the injected solution concentration, based on the average size of the vitreous space. Following the injections, Prednisolone 1%/ Gentamicin 0.3% ophthalmic ointment was applied topically to the eyes. The animals were placed on a heating pad until awake and monitored periodically for any signs of distress or complications.

### Optic Nerve Crush (ONC)

ONC was performed on 1-month-old C57BL/6J wild-type mice as in prior studies.^44,45^ Mice were anesthetized systemically with a mixture of xylazine and ketamine administered intraperitoneally, with 0.5% proparacaine eye drops applied topically. Under a dissecting microscope, the conjunctiva was lifted with fine forceps and cut using scissors, exposing the sclera. Forceps were used to manipulate and retract orbital fat and muscles, allowing for exposure of the optic nerve. Injury to the optic nerve was induced using curved fine-tip forceps to induce a focal crush injury to the optic nerve ∼1 to 2 mm behind the globe. Maximal pressure was used to close the forceps for 1 s. Use of fine-tip forceps facilitated avoidance of orbital vessels to prevent ocular ischemia. Bleeding during the procedure was categorized as none, minimal, moderate, or large. Mice were excluded for more than moderate bleeding. In each experiment, the ONC was performed in one eye only.

### Intravitreal injections of hiPSC-RGC

2-month-old wild-type C57BL/6J mice were anesthetized with a mixture of xylazine and ketamine administered intraperitoneally, with 0.5% proparacaine eye drops applied topically. For our injections, 2 µL of 1 X 10^6^ of hiPSC-RGCs transduced with an AAV2.7m8 virus-containing SNCG-eGFP vector was injected into the vitreous space of a C57BL/6 mouse’s eye (8-week-old, *n*=10 per mouse model) using a 33-gauge 0.5” needle attached to a 5 μL Hamilton syringe (Hamilton Company, catalog #: 7803-05, Reno, NV, USA). The contralateral eye was injected with 2 µL of 1X PBS, which served as an experimental control. Following the injections, Prednisolone 1%/ Gentamicin 0.3% ophthalmic ointment was applied topically to the eyes. The animals were placed on a heating pad until awake and monitored periodically for any signs of distress or complications.

### *In-vivo* ocular fundus imaging

Non-invasive retinal imaging was performed using Confocal Scanning Laser Ophthalmoscopy (“cSLO”, model Spectralis HRA, Heidelberg Engineering, Inc., Heidelberg, Germany) and Spectral-Domain Optical Coherence Tomography (“SD-OCT”, model Envisu R2200 UHR, Bioptigen, Inc., Morrisville, NC, USA) at 2-, 4-, and 6-weeks post-injection. Mice were anesthetized using Ketamine (100 mg/kg)/Xylazine (8 mg/kg), pupils dilated with a combination of 2.5% Phenylephrine/1% Tropicamide, followed by topical anesthesia to the cornea using 0.5% Proparacaine. Ocular protection against evaporative corneal desiccation during the imaging session was accomplished by using various combinations of artificial tears (Refresh, Genteal, and Balanced Salt Solution) and ocular eye shields. First, Wide-field (WF-55° FOV) and Ultra-wide field (UWF-105° FOV) cSLO images were collected in each eye with the plane of focus trained on the retinal ganglion cell layer/nerve fiber layer region. BluePeak Autofluorescence (BAF-cSLO, 486nm excitation/500-680 nm emission) images were collected with the optic nerve centrally positioned within the image field of view (FOV) frame. SD-OCT was performed last to assess the posterior pole retina and vitreous body structure. Posterior pole images included the posterior lens, vitreous body, retina, and choroid from a 50° FOV with the optic nerve centrally positioned. Following the imaging assessment, mouse eyes were covered with unmedicated ophthalmic ointment (Puralube Vet Ointment) and placed on a warmed heating pad (model 50-7220-F, Harvard Apparatus, Inc.) until recovered.

### Retinal histology and RGC quantification

A. **Flattened whole mount:** Eyes were harvested, fixed in 4% PFA at RT for 1 hour, and washed with 1X PBS, followed by isolation and dissection of the retina. Retinas were prepared as flattened whole mounts (*n*=6), washed with 1X PBS three times, and permeabilized in 0.5% Triton X-100 in PBS by freezing at - 80 °C for 15 minutes. Retinas were incubated overnight at 4 °C in a humidified chamber with either BRN3 (Santa Cruz, catalog #: sc-6026, Dallas, TX, USA), RBPMS (Sigma-Aldrich, catalog #: ABN1362, St. Louis, MO, USA), Tubulin β 3 (TUJ1; BioLegend, catalog #: 801201, San Diego, CA, USA), GFP (Novus Biologicals, catalog #: NB600-308, Littleton, CO, USA), Syntaxin-1 (abcam, catalog #:ab3265, Cambridge, UK), PSD95 (Cell Signaling Technology, catalog #: 3409S, Danvers, MA, USA), VGLUT2 (Cell Signaling Technology, catalog #: 14487S, Danvers, MA, USA), Iba1 (Novus Biologicals, catalog #: NB100-1028, Littleton, CO, USA), Ku80 (Cell Signaling Technology, catalog #: 2180S, Danvers, MA, USA), or Human Nuclear Antigen (Novus Biologicals, catalog #: NBP2-34342, Littleton, CO, USA). The next day, the primary antibody was removed, and the retinas were washed with 1X PBS five times for 5 minutes each, followed by incubation with appropriate secondary antibodies: anti-goat Alexa Fluor-594 (ThermoFisher Scientific, catalog #: A11080, Waltham, MA, USA), anti-mouse Alexa Fluor-594 (ThermoFisher Scientific, catalog #: A11020, Waltham, MA, USA), anti-rabbit Alexa Fluor-594 (ThermoFisher Scientific, catalog #: A21442, Waltham, MA, USA), and anti-rabbit Alexa Fluor-488 (ThermoFisher Scientific, catalog #: A11034, Waltham, MA, USA) for 1 hour at RT. The secondary antibody was discarded, and the retinas were washed with 1X PBS seven times for 5 minutes each. In the last step, retinas were incubated with DAPI (1 mg/ ml) (ThermoFisher Scientific, catalog #: 62248, Waltham, MA, USA) for 10 minutes at RT. Following incubation, the retina was washed with 1X PBS three times for 5 minutes each, and coverslips were then mounted on the slide using fluorescent mounting media (KPL, catalog #: 95059-434). Slides were observed under the Olympus FV1000 Confocal laser scanning microscope, and images were captured using 40X and 60X objectives with the use of appropriate filters and lasers. Images were analyzed using Fiji software.
B. **Cryosections:** Enucleated eyes were fixed in 4% PFA overnight at 4 °C. For cryo-sectioning, eyecups were dissected, and the lens was extracted (*n*=2). The retinal cups were first transferred to 15 % sucrose solution for 1 hour at 4 °C and were then again transferred to 30 % sucrose solution and incubated overnight at 4 °C. Eyes were embedded in optimal cutting temperature (OCT) compound, frozen on dry ice blocks, and stored at -20 °C for long-term storage. Using a cryostat, 14 μm sections of the retinal cups were generated. For immunohistochemistry, the cryo-sections were washed with 1X PBS three times for 5 minutes each. The slides were incubated with a blocking buffer (1 % BSA, 0.1 % Triton X-100, and 1 % normal donkey serum in 1 X PBS) in a humidified chamber for 1 hour at RT. The slides were then incubated overnight at 4 °C using the following antibodies: Ku80 (Cell Signaling Technology, catalog #: 2180S, Danvers, MA, USA) or Human Nuclear Antigen (Novus Biologicals, catalog #: NBP2-34342, Littleton, CO, USA). The next day, samples were washed with 1X PBS three times for 5 minutes each, followed by incubation with appropriate secondary antibodies: anti-mouse Alexa Fluor 594 (ThermoFisher Scientific, catalog #: A11020, Waltham, MA, USA) and anti-rabbit Alexa Fluor-594 (ThermoFisher Scientific, catalog #: A21442, Waltham, MA, USA) for 1 hour at RT. The coverslips were then mounted on slides using proLong Gold antifade reagent with DAPI (4′,6-diamidino-2-phenylindole) (Invitrogen, catalog #: P36935, Waltham, MA, USA). Slides were observed under an Olympus FV1000 Confocal laser scanning microscope, and images were captured using a 40X objective with the use of appropriate filters and lasers. Images were analyzed using Fiji software.

### Voltage reading of fluorescent GFP+ve cells and live two-photon imaging

Mice were euthanized by intraperitoneal injection of a lethal dose of ketamine (100 µg/g) and xylazine (10 µg/g). Eyes were enucleated and dissected in Ames’ medium (Sigma-Aldrich, St. Louis, MO, USA) continuously bubbled with 95% O_2_/5% CO_2_ under dim red illumination. Retinas were isolated, flat-mounted in a recording chamber, and perfused with oxygenated Ames’ medium. Bath temperature was maintained at 37°C using a TC-344B temperature controller (Warner Instruments, Hamden, CT, USA).

Patch pipettes were pulled from borosilicate glass capillaries (outer diameter 1.2 mm, inner diameter 0.69 mm; Sutter Instrument, Novato, CA, USA) using a P-97 micropipette puller. Pipettes were filled with an internal solution containing (in mM): 110 K-gluconate, 12 NaCl, 10 HEPES, and 1 EGTA (pH 7.4). CF633 (Biotium, Fremont, CA, USA) was added immediately before each experiment. Pipette positioning was performed using MPC-200 micromanipulators (Sutter Instrument) under infrared illumination using Kodak Wratten gelatin infrared filters (Eastman Kodak, Rochester, NY, USA) and a CCD-72 infrared camera (Dage-MTI, Michigan City, IN, USA). Electrophysiological recordings in cell-attached/voltage-clamp and/or whole-cell/current-clamp configurations were obtained using a PC-505B amplifier (Warner Instruments) and a Digidata 1440A digitizer controlled by Clampex software (Molecular Devices, San Jose, CA, USA).

Two-photon imaging was performed using a Fluoview FV1000MPE microscope (Olympus Corporation, Tokyo, Japan). Images were analyzed using ImageJ (National Institutes of Health, Bethesda, MD, USA), and electrophysiological and imaging data were further analyzed using custom MATLAB scripts (MathWorks, Natick, MA, USA). Action potentials were detected from high-pass-filtered traces (second-order Butterworth filter, 200-Hz cutoff frequency) using thresholds of -4 SD in cell-attached recordings and +4 SD in whole-cell recordings. Spike firing rates were calculated using 200-ms bins. All membrane potentials are reported without correction for the liquid junction potential. Corrected values would be approximately 16 mV more negative.

For the Z-stack video, the two panels on the left display the transmitted-light channel and an overlay of the green and red fluorescence channels. The green/red overlay was used to distinguish eGFP-positive hiPSC-RGCs from background autofluorescence, which was typically visible on both channels and therefore appeared yellowish. Because isolated retinas cannot be mounted perfectly flat for live imaging, different regions of the field of view may lie at different depths relative to the inner retinal surface. The panel on the right side of each video illustrates the position of selected image regions relative to retinal layers. Colored markers identify specific locations within the image, and each marker is associated with a cursor of the same color indicating the current depth position of that location relative to the retinal layer boundaries. Vertical offsets between cursors reflect local tilt of the mounted retinal preparation. Closely clustered cursors or a single cursor indicates minimal tilt.

### Optokinetic Response

The optokinetic response (OKR) was used to evaluate visual function every three months for up to a year after hiPSC-RGC injections in NMDA-treated mice. The OKR was assessed using OptoMotry software and equipment (Cerebral Mechanics, Inc., Medicine Hat, AB, Canada) as described in previous studies.^46,47^ Mice were placed on an elevated platform in the center of an arena consisting of four monitors with a camera above to image mouse behavior. The monitors displayed sine-wave gratings that rotated in a circular motion and varied in spatial frequencies, and the OKR was measured by determining the highest frequency at which mice can track a 100% contrast grid with head movements in the direction of the gratings. Measurements were performed by an investigator masked to the experimental treatments. Data are reported in units of cycles/degree.

### Statistics

Optokinetic responses over time were compared using analysis of variance (ANOVA) followed by Tukey post hoc comparisons for each treatment group. RGC counts and OKR final scores were compared using one-way ANOVA with Tukey post hoc comparisons between each group. All statistical analyses were calculated using Prism (GraphPad Software, San Diego, CA). All data are expressed as means ± standard error of the mean (SEM). *P*-values less than 0.05 were considered statistically significant.

## Author contributions

Conceptualization: V.V., V.R.M.C., and J.M.O.; methodology: V.V., V.R.M.C, and J.M.O.; validation: V.V., V.R.M.C, and J.M.O.; formal analysis: V.V., V.R.M.C., and J.M.O.; investigation: V.V., S.N., B.A.B., K.E.D., Y.L., R.S.K., K.S.S., A.G.R.; resources: V.R.M.C. and J.M.O.; data curation: V.V., S.N., and J.M.O.; writing – original draft: V.V; writing – review & editing: all authors; visualization: V.V., V.R.M.C., and J.M.O.; supervision: V.R.M.C. and J.M.O.; project administration: V.V., V.R.M.C., and J.M.O.; funding acquisition: V.R.M.C. and J.M.O.

## Acknowledgements

Rebecca J. Salowe helped review and edit the manuscript.

