## Supplemental Figures for "Re-evaluating donor-cell identity after hiPSC-RGC transplantation in mouse models of optic neuropathy"

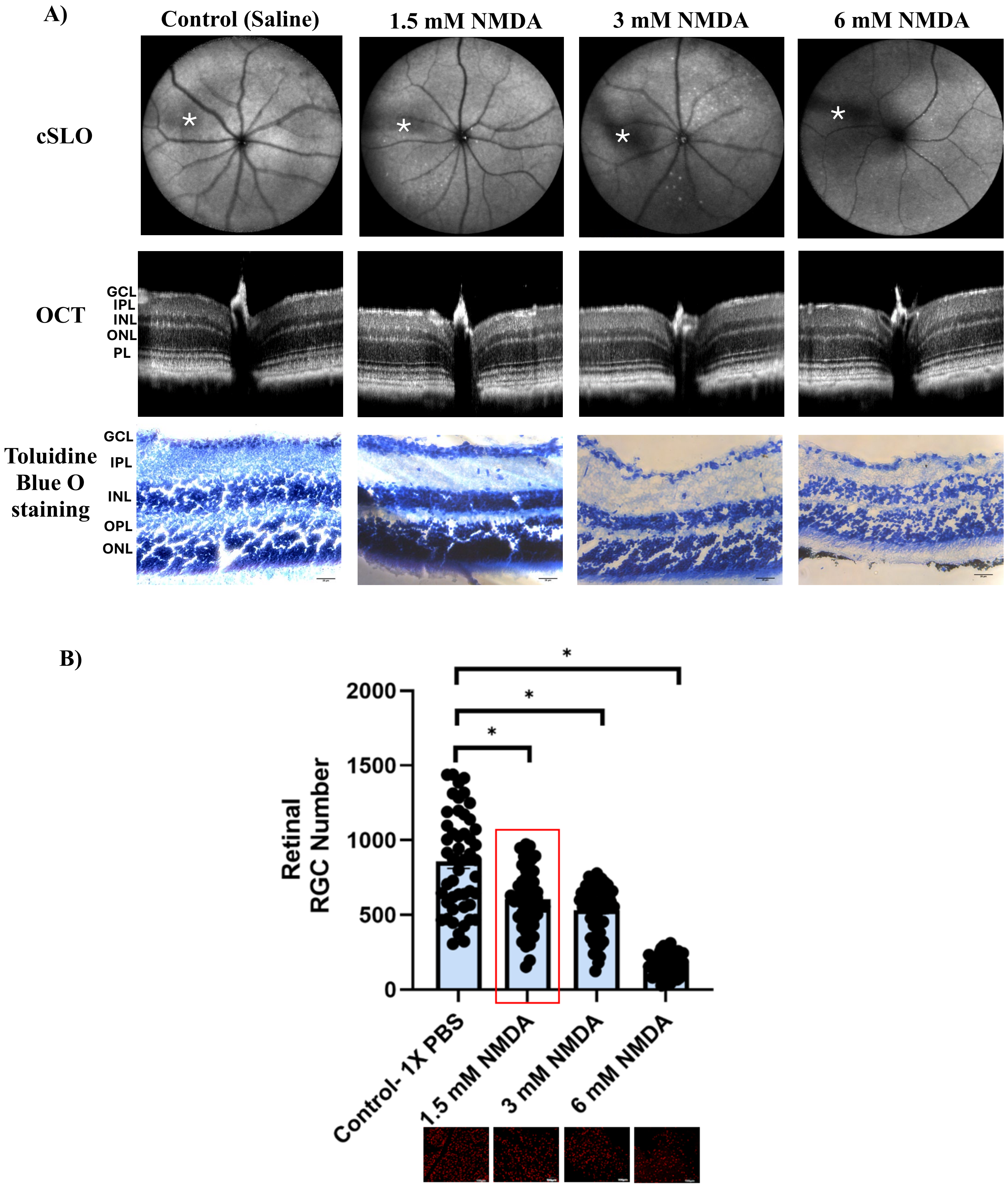


**Figure S1: Standardization of NMDA-mediated ganglion cell loss mouse model.** C57BL/6J wild-type mice were intravitreally injected with 1X PBS or with NMDA at 1.5 mM, 3 mM, or 6 mM (*n*=5/group). One month after treatment, we imaged the mice's eyes using A) cSLO, SD-OCT, and toluidine blue O staining (images were taken at 40X magnification). B) Quantitative representation of the number of Brn3$^{+ve}$ RGCs in the retina among all groups of NMDA-treated mice. * Indicative of intravitreal injection track.

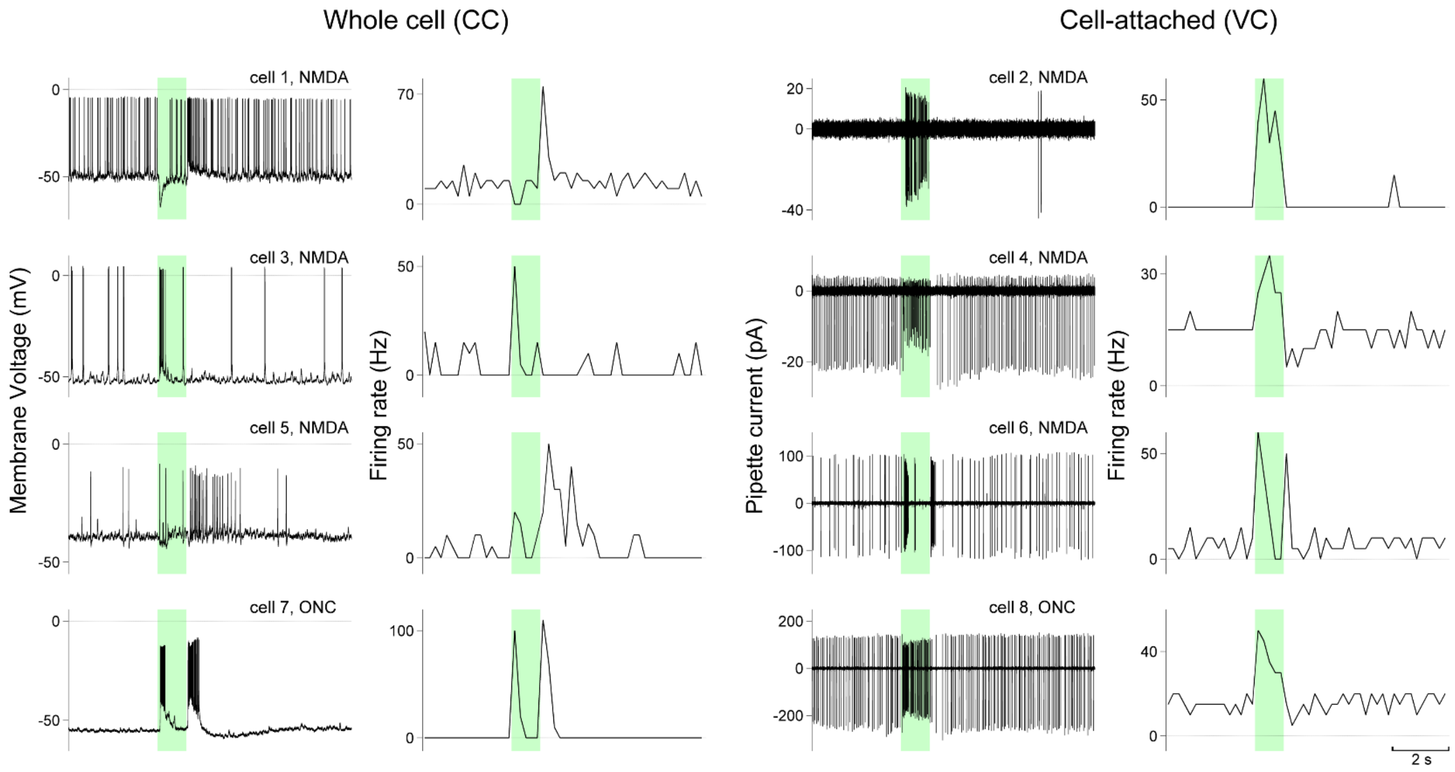


**Figure S2: Light responses of eight GFP$^{+ve}$ cells integrated into the mouse retina.** Patch-clamp recordings obtained in either whole-cell/current-clamp or cell-attached/voltage-clamp configurations were used to monitor light-evoked spike firing activity. For each cell, the left panel shows membrane potential (whole-cell/current clamp) or high-pass-filtered pipette current (cell-attached/voltage clamp), and the right panel shows spike firing rate calculated using 200-ms bins. Sharp vertical deflections in the membrane potential and pipette current traces represent action potentials. Colored bars indicate periods of light stimulation. The light intensity was $4 \times 10^5$ photons μm$^{-2}$ s$^{-1}$ for all cells except cells 3 and 7, for which intensities of $1 \times 10^5$ and $2 \times 10^5$ photons μm$^{-2}$ s$^{-1}$ were used, respectively. Membrane potentials were not corrected for the liquid junction potential; corrected values would be approximately 16 mV more negative. Cell 1 is the same as shown in Figure 3.

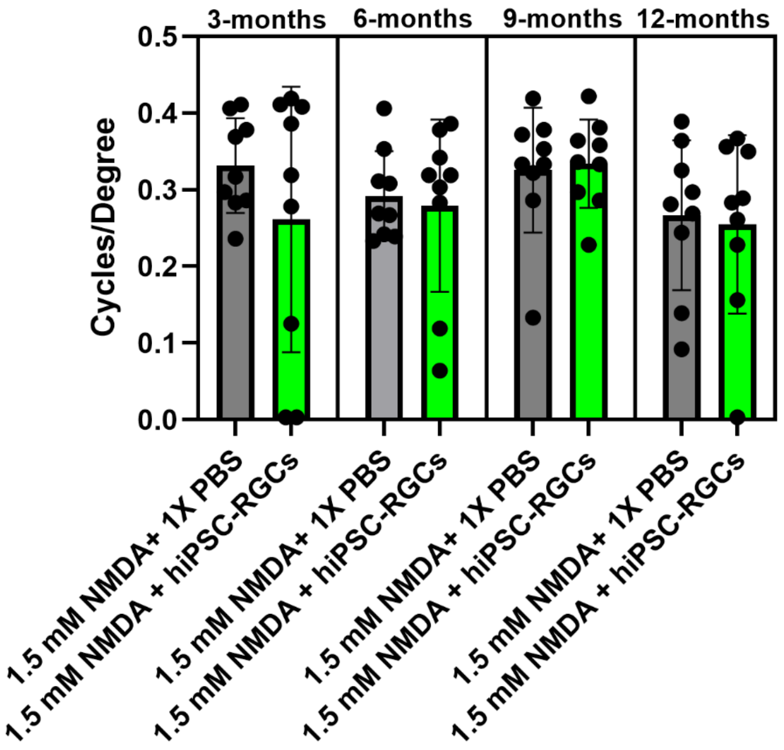


**Figure S3: OKR responses documented to detect improvement in visual acuity.** OKR scores compared between the right eyes treated with 1.5 mM NMDA and hiPSC-RGC injection and the contralateral eyes injected with 1.5 mM NMDA and 1X PBS (*n*=9). OKR scores did not differ significantly between the two treatment groups at any time point over the year. 0.3-0.4= Normal vision, whereas 0 means visually impaired.

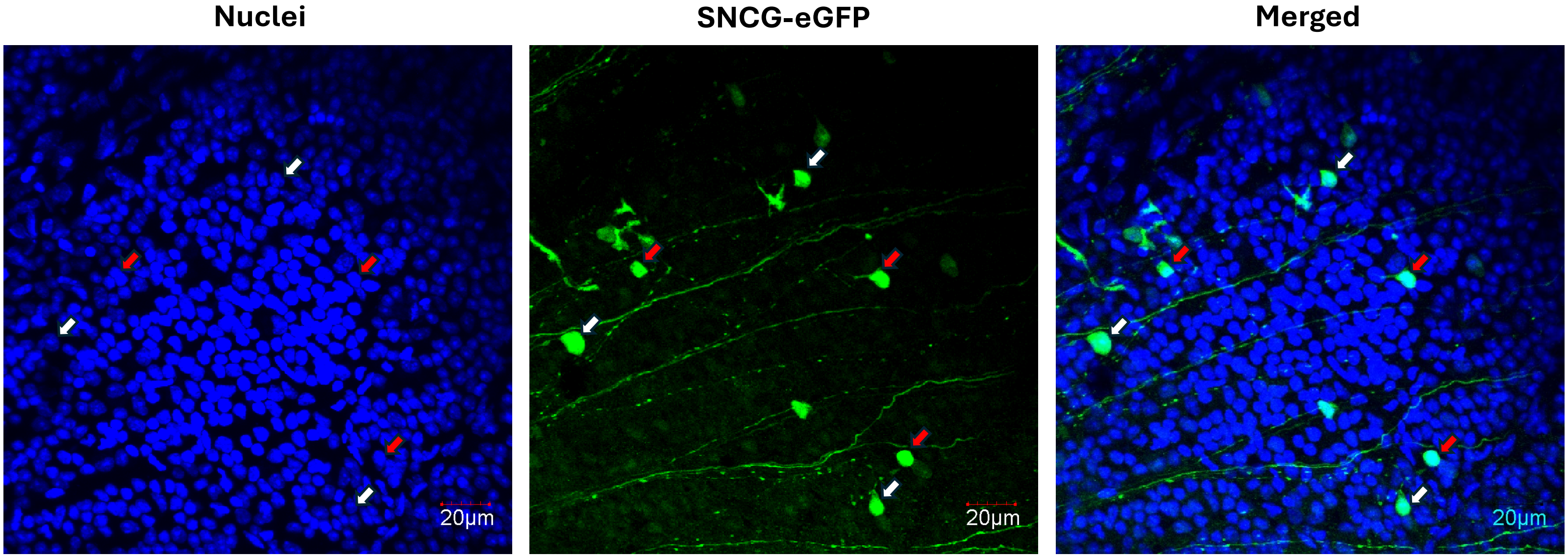


**Figure S4: Flat-mount image of *SNCG*-GFP$^{+ve}$ cells detected in the mouse retina one year after transplantation.** Red arrows point to the nuclei of cells with homogeneous DAPI staining, indicative of human cells. White arrows point to the nuclei of cells with prominent nucleoli, indicative of mouse cells.

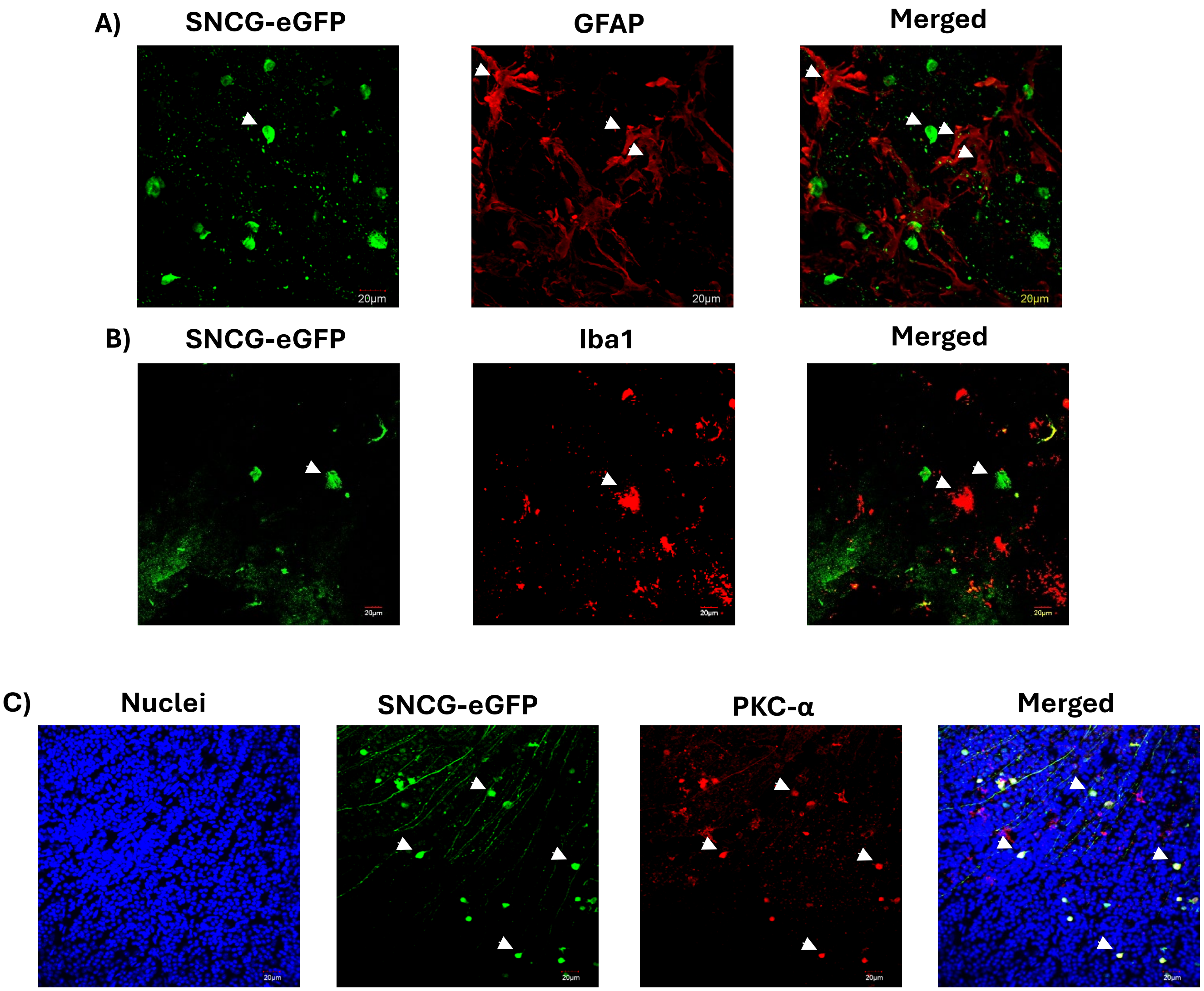


**Figure S5: Mouse retinas transplanted with hiPSC-RGCs co-stained for other retinal cell markers.** Flattened whole-mount images of GFP$^{+ve}$ cells co-stained for the following markers: (A) GFAP, (B) Iba1, and (C) PKC-α, indicative of Müller glia, microglia, and bipolar cells, respectively.
